# A Standardized Protocol for Quantifying the Behavioral Dynamics of Food-seeking in Mice

**DOI:** 10.1101/2021.10.09.461623

**Authors:** Dong-Soo Ha, Young Hee Lee, Kyu Sik Kim, You Bin Kim, Hyung Jin Choi

## Abstract

Food is generally hidden in a natural environment and require free-living animals to search for it. Although such food-seeking behaviors involve motivation and exploration, previous studies examined food-seeking simply by measuring the time spent in the food zone or the frequency of pursuing food-cued context. Moreover, after discovering food, animals need to taste and smell it in order to evaluate their nutritional value or possible toxicity. However, researchers could not easily distinguish food-seeking from food-evaluating behaviors because food was visible or accessible throughout each test. Herein, we describe a behavioral protocol that triggers animals to show the behavioral dynamics of food-seeking (e.g., navigation, nose-digging, and paw-digging) and that exclusively elicits food-seeking without provoking any other food-evaluating behaviors. First, we prepared an open-field box with the floor covered with bedding. After we hid foods under the bedding of each corner, the test mice were habituated in this arena for four days (pre-test phase). On the next day (test phase), they were placed under the same conditions, but the foods previously hidden were removed. This process enabled the mice to perceive their surroundings as a food-hidden environment, which induced the animal to exhibit sustained food-seeking. In conclusion, the protocol presented here is a powerful method for provoking multiple forms of food-seeking and quantifies food-seeking independently from other food-related behavioral stages.

## INTRODUCTION

Nature comprises a wide range of nutritious resources. However, these are generally hidden from animals and cannot be freely obtainable whenever needed. Thus, to secure energy resources essential for survival, animal neural systems have evolved to direct food-seeking behaviors (REF 1, 2). Furthermore, as animals acclimatize to their living environment, the location of the food can be anticipated. Thus, based on their previous experiences, animals may move directly to areas where food could be found.

Food-seeking is not a single-step process. Firstly, animals start moving around their environment with the motivation for food. Then, as soon as they locate a specific area where food could possibly be found, animals actively search food with their paws and nose. Likewise, animals in natural environments exhibit food-seeking behavior in a multi-step procedure.

Moreover, food-seeking is followed by other food-related behaviors because discovering food does not necessarily mean that animals consume it. As animals in nature did not encounter such food before, they could not immediately confirm that the item they found was edible. Thus, animals need to taste, smell, and inspect their discovery to evaluate their nutritional value and possible toxicity. After finishing the food-evaluating behavior, animals decide whether they should consume it. (REF 3)

Provoking food-seeking increases the likelihood of animals exhibiting food-evaluating behavior, as these are sequential behaviors. Similarly, activating food-evaluating behavior can increase the possibility of animals consuming food. Owing to such connections, the exclusive measurement of each behavior is challenging. Herein, we introduce a behavior test that specifically targets food-seeking behavior without provoking any other behavioral stages. Ultimately, our protocol provides a powerful foundation for scrutinizing feeding behaviors in a more natural and efficient way.

### Comparison with other methods

Although food-seeking is a multi-step process, previous studies have investigated a food-seeking as a single-step behavior. For instance, “latency to enter food zone” or “time spent in food zone” was regarded as an equivalent to food-seeking (REF 4). In this study, a shorter latency indicated a more significant motivation for food. However, they did not show any evidence that mice arrived in the food zone earlier because they wanted food. Furthermore, the time spent in food zone also includes non-food-related behaviors such as grooming and sleeping. Therefore, this previous method could be an index related to food-seeking, but it lacks specificity and does not directly measure food-seeking.

Moreover, the term “seeking” can be only used when targets are not identifiable. However, as the identifiable foods are consistently visible during the food-zone test, researchers could not find the exact start- and end point of food-seeking. Thus, behavioral indices that rely only on food zones cannot correctly show the motivation for food-seeking while having a high risk of entailing other behavioral modalities.

Furthermore, various studies regarded the “number of pressing food-cued levers” during the operant conditioning test as food-seeking (REF 5). However, the task did not require animals to navigate or move around their surroundings. Moreover, lever-pressing is not a natural form of food-seeking shown by free-living animals. Hence, the operant conditioning task only measures the intensity of motivation and fails to elicit the multiple phases of food-seeking demonstrated in free-living animals.

### Development of the protocol

Conventional methods were unable to measure the exact moment of food-seeking. In addition, they could not elicit multiple stages of food-seeking. To overcome these limitations, we established a behavioral protocol that induces multiple stages of food-seeking without triggering food-evaluating behaviors. We first divided food-seeking procedures exhibited by free-living animals into (1) navigation, (2) entry to food-associated areas, and (3) digging.

As soon as mice encounter an environment different from their home, they move around their surroundings to learn where they are. In addition, they navigate through the arena to pursue specific goals, such as food. We call this stage “navigation”. To allow mice to navigate, we used an open-field box, which is a widely used chamber in animal behavior research.

Moreover, the involvement of external cues can promote food-seeking behavior. Animals seek more actively when they associate food with previously neutral cues. In addition, this association elicits an adaptive motor behavior for food. (REF 6, 7) Thus, we filled the open-field box with beddings and hid food at the beddings of each corner. We hypothesized that associating geographical cues with food could induce animals to seek food more efficiently.

Furthermore, among various forms of food-seeking, digging is widely known as a naturalistic behavior when mice forage for food. (REF 8) Repetitive exposure to the same condition allows mice to learn that they will find food when they dig the beddings of each corner (pre-test phase). However, during the test phase, we placed the mice under the same conditions without food under the bedding. The time spent digging the food-associated areas indicated the amount of food-seeking behaviors displayed by the mice. Since there were no foods that could be found throughout the test phase, we were able to measure food-seeking without provoking any food-evaluating behavior. In addition, no food means that animals will exhibit sustained food-seeking throughout the entire test duration, which provides sufficient data for quantification.

### Experimental design

Mice were conditioned in a white acrylic open field box (32cm × 32cm × 30cm) covered with wood shaving bedding, where food was hidden under the bedding at each corner. The thickness of the stacked bedding was approximately 1.5 cm. Conditioning was held in this arena seven times (twice per day, each in the morning and afternoon, 10 minutes per trial). We called this part of the test the “pre-test phase.” After conditioning, the mice were placed in the same situation, but without food hidden the following day. We call this part the “test phase”. In the case of a cross-over study, to eliminate the memory of the previous context, the mice were re-conditioned to the food-hidden environment three times (two days after the test phase). Herein, conditioning (pre-test phase 2) was performed separately for two days. The next day, the mice were again placed in an identical situation, but without hidden food (test phase 2).

Previous exposure to the food-hidden environment allows mice to learn that food is placed under the bedding of each corner. However, the absence of foods during the test phases deceived the mice, as they still thought that foods should be hidden in the arena. Digging duration, distance moved, frequency of visiting the corners (food-associated area), latency to start digging, first minute velocity, and total velocity were analyzed to quantify food-seeking.

Wood shaved beddings were used in this protocol for two reasons. First, they are the same material that home cages are filled with, which will reduce the anxiogenic effect during the experiment. Furthermore, this makes it more challenging for mice to seek food. When mice dig the beddings of a certain area, beddings from other areas are likely to fill in. This makes it difficult for animals to conclude that foods that they are looking for are removed. Ultimately, it induces sustained food-seeking in mice.

### Applications of the method

#### Foods and objects

Animals not only seek foods, but also seek novel objects. Thus, modifying environments and substituting food for other objects such as water, novel objects, females or males, and warm or cool substances could expand the goal of the behavior test. Anything that can motivate the animal to seek can be utilized with this protocol. Anything that can motivate the animal to seek can be utilized with this protocol. In addition, we observed mice digging the acrylic floor even though the bedding was cleared. Such obsessive seeking behavior can be used as an index to examine the behaviors of obsessive-compulsive disorders (OCD) or anxiety mice models.

#### Maze

Repetitive exposure to the food-hidden environment allows mice to expect food when they are reintroduced to that environment. However, we removed the food to deceive the mice. This procedure successfully evoked sustained food-seeking because we previously conditioned them to think that foods should be hidden in the arena. Researchers can broadly apply the psychological concept of deception to any other behavioral maze, in order to measure the “seeking” behavior in animals.

#### Studying neural networks

This protocol suggests a novel approach for feeding behavior research. It provides a pioneering foundation for investigating the neural network of each feeding behavior, as we isolated certain stages of these behaviors. Moreover, artificial neural manipulation tools such as optogenetics and chemogenetics can be applied to this protocol, thereby unraveling the function of specific neural pathways for food-seeking. Moreover, substituting food to objects would allow researchers to study the neural mechanisms of other contexts, such as novelty-seeking.

### Limitations

Digging does not always imply food-seeking because it is also a representative behavior of compulsivity. Thus, to verify that mice dig to seek food, this protocol could be paired with compulsivity tests such as the marble burying test. Moreover, mice generally show anxiety in the central zone of the arena. Therefore, they tend to stay longer at the corners or edges of the arena. Thus, researchers may change the food-associated areas to the center of the arena to provide a more robust version of food-seeking.

## MATERIALS

### Biological materials

#### Mice

10-week old C57BL/6 and C57BL/6J mice were used. The mice were kept in a room at a temperature of 22 ± 1 °C. On a 12-hour light-dark cycle, the light cycle was 8 am-8 pm and the dark cycle was 8 pm-8 am. If mice had to be conditioned for chronic deprivation, they were allocated into separate cages. During the test phase, the mice were kept ad libitum. Mice were placed in a room to stand with optimal temperature and light before experimenting in a sound-proof behavioral chamber. The experiment was maintained at a temperature of 22 ± 1 °C. All procedures were approved by the Institutional Animal Care and Use Committee of Seoul National University. **\*Caution** Mice that are younger or older can be used (6-week~1-year), but for optimal experimental conditions, 8~10weeks w recommended. The test phase was performed in a light cycle.

#### Reagents

Reagents are required only when chemogenetic methods are used. AAV8-hSyn-DIO-hM3Dq-mCherry virus (Addgene, Plasmid #50475, titer 1.9×10^13). was injected into LH Leptin receptor-expressing (LepR) neurons of LH LepR Cre mice. Clozapine-n-oxide (Sigma, 34233-69-7, CNO, 1 mg/kg, I.P.) and saline (1 mg/kg, I.P.) were injected during the test phase.

### Equipment

#### Recordings

Cameras with no distortion (ELP-USB4KHDR01-KV100, Ailipu Technology Co., Ltd; Microsoft Lifecam HD 3000, Microsoft) were used for the behavioral analysis. The quality of the recording was 720X480 for ELP and 1280X1080 for Microsoft Camera.

#### Open-field box

A white acrylic box with a size of 32cm × 32cm was used. The thickness of the box was 1cm, making the area of the inner space 30cm × 30cm.

#### Beddings

All bedding used in the experiments was from KOATECH Aspen Bedding, which was autoclaved before the experiment.

## PROCEDURE

### Habituation

1. Handle the test animals for at least 5 consecutive days (for 3-10 minutes per day) before starting the assay. **\* CAUTION** The test animals should have been living in a similar environment and have similar body weight since these factors can affect the activity of the animal. Every step was completed before 6 PM. The duration of the handling process can be modified based on the animals’ stability.
2. The test mice were placed into the sound-proof behavioral chamber and allowed to habituate for at least 30 min. The chamber was set to the test conditions. **\* CAUTION** Light shed in the arena should not induce anxiety-like behaviors in mice. The light intensity was measured at the center of the open-field box.
3. Arena habituation would be ideal, but this step could be ignored because of the repetitive conditioning steps that will be performed later.

### Preparation for the pre-test phase (Day 1)

1. Prepare a white acrylic open field box.
2. Place two raisins (or chocolate-flavored crackers) at each corner of the open-field box. The number or amount of food can be modified freely.
3. Fill the floor of the box with wood shaving beddings, with the thickness of the stacked beddings being approximately 1.5 cm. Make sure to hide the foods with the beddings so that the foods are not visible. **\* CRITICAL** Stacking too many beddings will be a difficult task for mice and may induce anxiogenic effects, while a small amount of beddings enables foods to be exposed even before the start of digging. Moreover, beddings should be distributed evenly throughout the surface of the arena.

### Pre-test phase (Day 1-4)

1. To make mice conditioned to the settings, place one test animal at the center of the arena and close the door of the behavior chamber. The mouse were allowed to explore for 10 min.
2. Repeat the procedures above for every mouse. **\* CRITICAL** Open-field box was cleaned with 70% EtOH and distilled water after each trial. For every trial, fresh beddings and food were used.
3. Repeat procedure 1 and 2 for at least seven times (twice per day, each in the morning and afternoon, 10 min per trial).

### Preparation for the test phase (Day 5)

1. Prepare the identical open field box used in previous sessions.
2. The floor of the box was filled with beddings, with the thickness of the stacked beddings being approximately 1.5 cm. Make sure that there is no food under the bedding. **\* CRITICAL** Again, do not stack too much bedding. Beddings could be reused within the trials of the same mouse if the feces were removed in each trial.

### Test phase (Day 5)

1. Place one test animal at the center of the arena and close the door of the behavior chamber. The mice were allowed to explore for 10 min.
2. Repeat the procedures described above for every mouse. **\* CRITICAL** The open-field box was cleaned with 70% EtOH and distilled water after each trial. Make sure that every mouse encounters fresh bedding.

### In case of a cross-over study (Day 8-9) – Pre-test phase

1. After two days (rest the mice on day 6-7), in order to diminish the memory from the test phase, repeat the procedures depicted in the “Pre-test phase”. The number of trials was reduced half (**Table 1**). **\* CAUTION** Make sure that foods are hidden under the bedding.
2. The mice were conditioned to the food-hidden environment three times (two days after the to-be-analyzed test). Conditioning should be performed separately for two days.
3. Repeat procedure 1 and 2 equally for every mouse. **\* CRITICAL** The open-field box was cleaned with 70% EtOH and distilled water after each trial. Make sure that every mouse encounters fresh bedding.

**Table 1.**
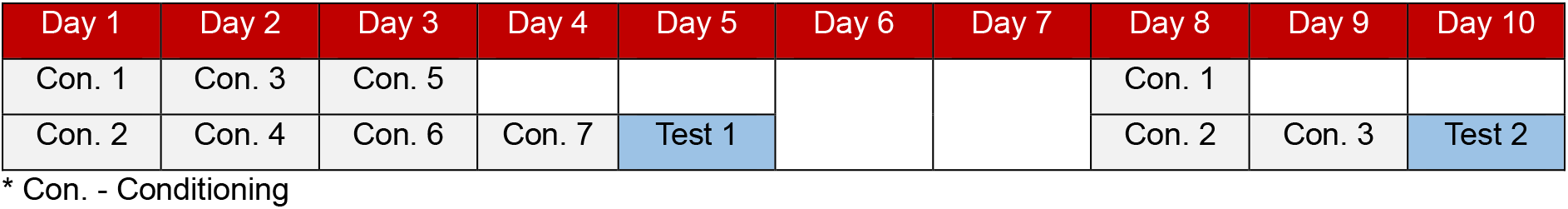
Timeline of the food-seeking test

### In case of a cross-over study (Day 10) – Test phase

1. The mice were again placed in an identical situation, but without hiding the food. **\* CAUTION** Make sure that the food is removed at this time.
2. Repeat procedure 1 and 2 equally for every mouse. **\* CRITICAL** The open-field box was cleaned with 70% EtOH and distilled water after each trial. Make sure that every mouse encounters fresh bedding.

### Video recordings

Conditioning and testing were performed under constant light conditions, whereas recordings were performed above the center point of the arena.

### Automatic analysis

Recorded videos were reviewed and analyzed using EthoVision XT (Noldus). Food-seeking indices such as time spent in food-associated areas, total velocity, first-minute velocity, entry to food-associated areas, latency to start food-seeking, and consecutive visits to the food-associated area were quantified. The time spent in food-associated areas included the animal’s motivation for food. The total velocity reflects the animal’s motivation for food. In addition, the first-minute velocity better represents food-seeking because their motivation for food would be the highest as soon as the animals are placed in the arena. Entry to food-associated areas indicates the frequency of the mouse visiting each food-associated area. The latency to start food-seeking indicates motivation for food. Finally, a consecutive visit to the food-associated area indicates the frequency of the mouse visiting each food-associated area in a clockwise or counter-clockwise order, reflecting the navigation phase of food-seeking. Animals in the arena were traced by body points for 15 min. Food-associated areas (corners) were designated as 1/16 (or 1/9) areas of the arena. Animal nose point detection could be more useful when analyzing animals’ behavioral directions.

### Manual analysis

An Observer XT (Noldus) was recommended. Digging behavior was analyzed manually and categorized into three subcategories: 1. Digging with nose, 2. Digging with paws, 3. Digging even though the bedding was cleared. The criterion for each behavior was as follows: First, nose-digging was counted when mice pushed their nose into the bedding. However, this was restricted to the event only when the bedding was moved because of this behavior. Second, digging with paws was counted when mice used their paws to dislocate bedding and find food. The last behavior was counted only when the mice kept searching the acrylic box even though the bedding had been cleared to show that there was no food. “Cleared bedding” was defined when there were no beddings in front of the faces of the mice. Each behavior was counted only if it was more than 0.2 seconds, since less than 0.2 seconds could include errors lasting than its duration. The time spent digging in each zone was combined to determine the total duration of digging.

## TROUBLE SHOOTING

### Size of the food-associated area

The size of the food-associated area (1/9 or 1/16 of the arena) was determined based on the size of the mouse. However, this does not necessarily mean that it is the exact area that animals perceive as a food-associated area. Thus, researchers can designate an area based on the size of the animal or food.

### Lighting

The lighting of the behavior chamber may affect general activities and food-seeking. Increased anxiety due to high light intensity may reduce overall locomotion. Furthermore, when light is unevenly distributed to the arena, mice remain in a dimmer region. Therefore, an indirect light source that can minimize anxiety of the test animals is recommended. Moreover, when camera zoom-in is unavailable, a reduction in light intensity is highly recommended.

### Beddings

The bedding was evenly distributed across the surface of the arena. If not, the area with more bedding can be a potential location for nesting. In addition, it is essential to avoid food exposure on the surface.

### Number of cameras

A sufficient number of cameras is beneficial for behavior analysis. In our apparatus, we used two cameras to maximize the range of view in the mouse behavior analysis. The top view was for locomotive analysis, and the side view was used for additional analysis.

## RESULTS

Optogenetics and chemogenetics are widely used to manipulate specific types of neurons in the animal brain. In this protocol, we used both methods to activate the lateral hypothalamic (LH) leptin receptor-expressing (LepR) neurons, and monitored the change in food-seeking behaviors before and after the manipulation. Herein, we describe the results of chemogenetics.

We used the same protocol as described in the article. Mice were conditioned in an open field covered with wood shaving bedding, where food was hidden under the bedding at each corner. After conditioning, the mice were placed in the same situation, but without food hidden **(Figure 1)**. The time spent in food-associated areas (corners) and the field (non-corner areas), the velocity of mouse movement, and the velocity of mice in the first minute were automatically analyzed during the chemogenetic-manipulating method **(Figure 2A-C)**. Furthermore, the track visualization and heat map of the 10-min videos were generated **(Figure 2D)**. As a result, the activation of LH LepR neurons induced a shortened latency to start food-seeking, indicating that these neurons may be involved in accelerating the food-seeking process. The track visualization data showed that locomotion from one food-associated area to another was increased after activation. Moreover, the time spent food-seeking was analyzed using the analysis criteria described earlier, and this was significantly increased when the neurons were stimulated. **(Figure 3)** These results demonstrate that LH LepR neurons drive food-seeking behavior. In conclusion, these data together support that the protocol is a sufficient tool for analyzing food-seeking without inducing any subsequent feeding behaviors in mice.

**Figure 1.**
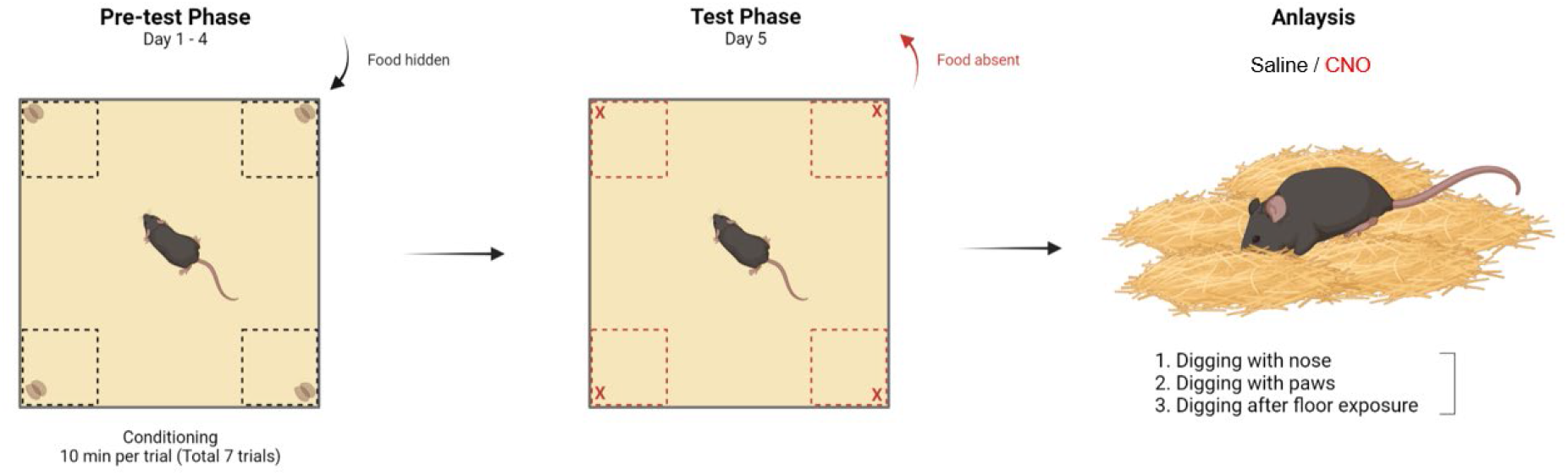
Graphical procedure of the food-seeking test. Mice were conditioned in an open field covered with wood shaving beddings, where food was hidden under the bedding at each corner. After conditioning, the mice were placed in the same situation but without food hidden.

**Figure 2.**
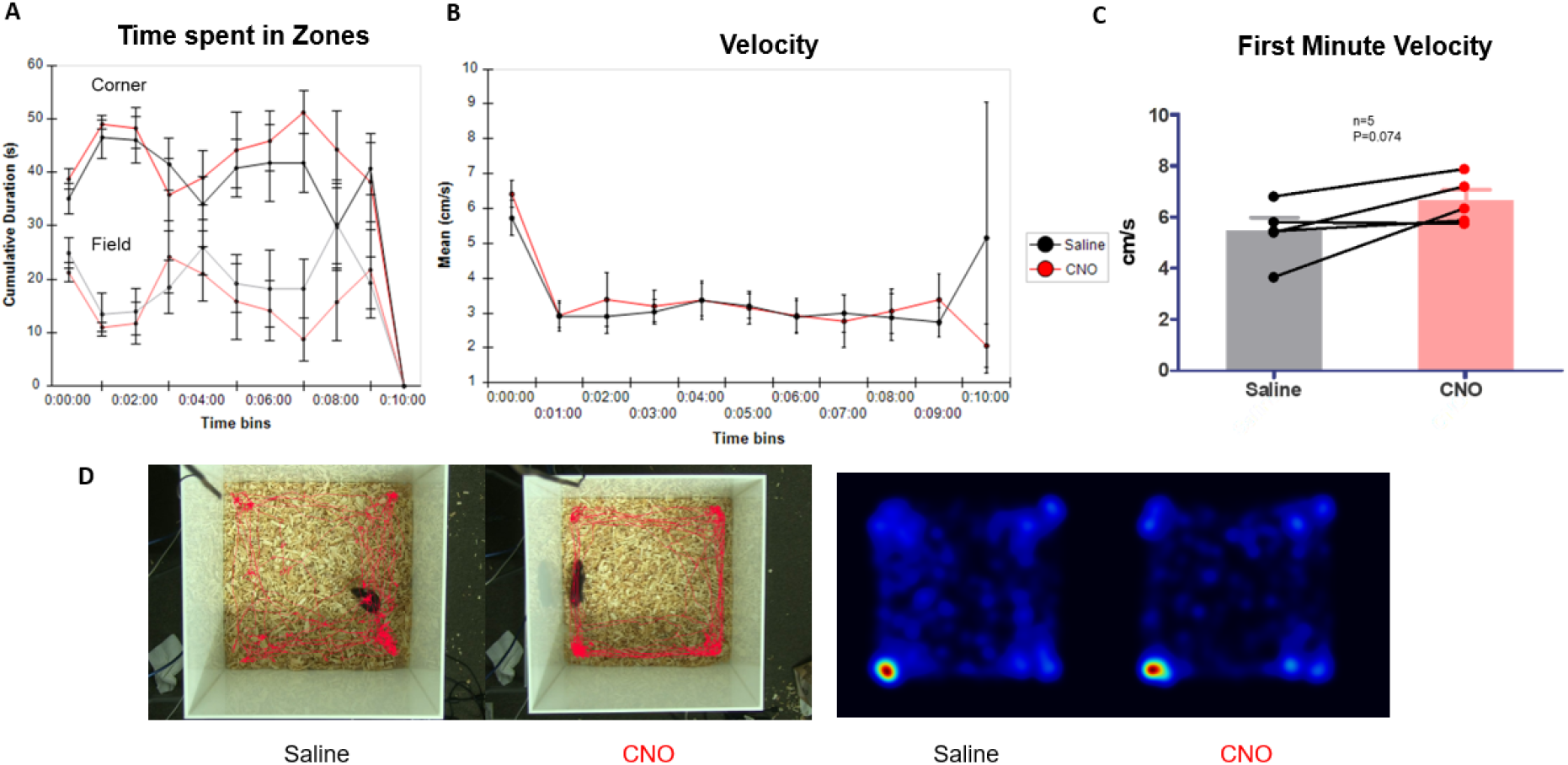
Automatic analysis of food-seeking during chemogenetic modulation. **A**. Time spent in the corners and field, **B**. Velocity of mice movement by time bins, **C**. Velocity of mice at the first minute, **D**. Track visualization and heat-map of the 10 min videos were analyzed.

**Figure 3.**
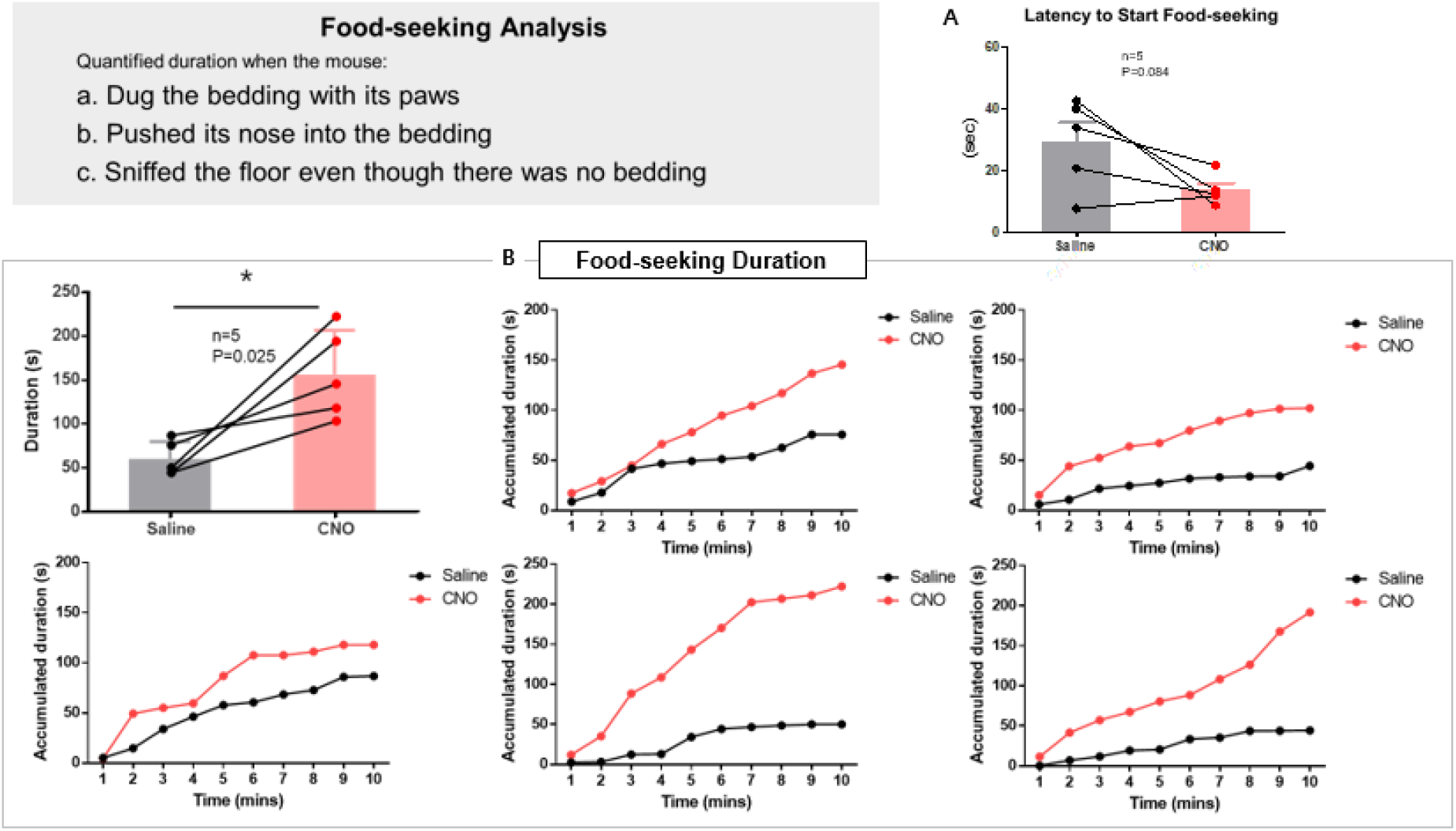
Quantification of food-seeking based on the analysis criteria. The time spent seeking food was analyzed using the quantification criteria. **A.** Latency to start food-seeking, **B.** Food-seeking duration. Individual data for each mouse were further analyzed using time bins that are displayed in the five graphs above.

## AUTHOR CONTRIBUTIONS STATEMENTS

D.S.H. developed the experiments and analytical framework, performed the experiments, analyzed data, and wrote the manuscript. Y.H.L. performed the experiments and analyzed data. K.S.K. performed the experiments, analyzed data, and partially contributed to writing the manuscript. Y.B.K. performed the experiments and analyzed data. H.J.C. supervised and modulated the entire study.

## REFERENCES

1. Roitman et al., Dopamine operates as a subsecond modulator of food seeking, J. Neurosci. 2004 February 11;24:1265–1271.

2. Wyvell et al., Intra-accumbens amphetamine increases the conditioned incentive salience of sucrose reward: Enhancement of reward ‘wanting’ without enhanced ‘liking’ or response reinforcement, J. Neurosci. 2000 November 1;20:8122–8130.

3. Matthias Laska, Freist Pamela, Krause Stephanie, Which senses play a role in nonhuman primate food selection? A comparison between squirrel monkeys and spider monkeys, Am. J. Primatol. 2007;69:282–294

4. Carus-Cadavieco et al., Gamma oscillations organize top-down signalling to hypothalamus and enable food seeking, Nature February 09 2017;542:232–236

5. Martín-García et al., New operant model of reinstatement of food-seeking behavior in mice, Psychopharmacol. (Berl.). 2011 May;215:49–70

6. Michael A. Cowley, … Robert V. Considine, in Endocrinology: Adult and Pediatric (7th Edition), 2016

7. Richard Mattes, Sze-Yen Tan, in Nutrition in the Prevention and Treatment of Disease (3rd Edition), 2013

8. Pond et al., Digging behavior discrimination test to probe burrowing and exploratory digging in male and female mice, J. Neurosci. Res. 2021 May;99:2046–2058

